# The role of sex-based developmental class in predicting group reproductive output in wild callitrichids

**DOI:** 10.1101/047969

**Authors:** Mrinalini Watsa, Gideon Erkenswick, Efstathia Robakis

## Abstract

Prior research on cooperative breeders has considered correlations between group reproductive output (GRO) and the number of individuals in each age-sex class, but without controlling for uneven sampling efforts, the underlying effects of group size, and pseudoreplication at the group and species levels. Among callitrichids, age-sex classes do not provide meaningful categories, as individuals within an age-sex class can demonstrate varying reproductive development due to reproductive dominance of a few individuals per group. This study re-assesses the drivers of GRO in callitrichids by a) conducting a meta-analysis of published studies of callitrichid group composition; b) determining a novel method to assign developmental class based on reproductive morphology; and c) utilizing a multistep modelling approach to assess whether any sex-based developmental class predicts both the presence and the numbers of surviving offspring among free-ranging saddleback (*Leontocebus weddelli*) and emperor tamarins (*Saguinus imperator*) in Peru. The meta-analysis utilizing a historical dataset revealed that adult females and group size, but not the number of adult males is significantly correlated with GRO. Statistical models of the new dataset revealed that only mature males predicted if groups had any infants at all, but that the number of surviving infants was predicted by mature females and group size. Thus mature males appear to be necessary for groups to raise any infants, but mature females and a larger group size increase group reproductive output overall.

## Introduction

Callitrichids exhibit a cooperative breeding system in which offspring receive care from non-biological helpers, or alloparents (Garber 1997; Jennions 1994; Sussman and Garber 1987). Groups typically consist of a single breeding female (although other females may be present) and variable numbers of adult and sub-adult males. All adults in a group participate in infant rearing, including infant provisioning and transportation (Bales et al. 2000; Garber et al. 1984; Goldizen and Terborgh 1986). Despite the monopoly of breeding by a single adult female in most cases, callitrichids are rarely monogamous, but rather display a range of flexible mating strategies even within groups (Garber 1997; Garber et al. 2015; Goldizen et al. 1996; Sussman and Garber 1987; Terborgh and Goldizen 1985).

One of the primary arguments for the presence of helpers, typically unrelated adult males or natal subadults, is that they alleviate the cost of rearing energetically expensive twin offspring that constitute over 80 % of all births in callitrichids (save *Callimico*) (Tardif 1997; Wislocki 1939). Alloparenting behaviors by helpers benefit offspring survival, and thus increase group reproductive output (GRO) (Bales et al. 2000; 2001; Boulton and Fletcher 2015; Garber 1997; Heymann 2000; Koenig 1995). By investing in the care of offspring, helpers could incur indirect fitness benefits if they are related to the biological parents; they also benefit the group by providing increased vigilance and protection from predators, or access to valuable resources (for reviews see Bales et al. 2000; French 1997; Tardif 1997). Prior research suggests that the effects of helpers on GRO are not uniform, and can vary based on helper sex and social status (Bales et al. 2000). There may also be species differences in how helpers of different age-sex classes influence GRO due to differing costs of infant-rearing between species (Díaz-Muñoz 2015; Heymann 2000).

Four cases in the published literature have attempted to explain variation in GRO by correlational analyses of group demographics. First, a study by Garber et al. (1984) found that the average number of infant moustached tamarins (*Saguinus mystax*) that survived to become juveniles was significantly positively correlated with the number of adult male helpers in a group. A follow up to this study further indicated that groups with one adult male had one third the number of dependent offspring that groups with three or more adult males did, independent of group size (Garber 1997). Second, a review of research on wild common marmosets (*Callithrix jacchus*) revealed that the number of juveniles was significantly correlated only with the number of adult males, and no other age-sex class (Koenig 1995). Third, using a large dataset on golden lion tamarins (*Leontopithecus rosalia*), Baker et al. (1993) calculated a higher mean number of offspring for two-male groups than in single-male groups, only including adult non-natal males in these analyses. Lastly, Bales et al. (2000) examined the effects of particular alloparents on GRO in the same population of *L. rosalia* by classifying alloparents in two ways, a) as “helpers”, defined as all animals over 18 mo of age other than the reproductive pair and reproductive subordinate females and b) “adult males”, both breeding and nonbreeding (Garber 1997; as with Garber et al. 1984; Koenig 1995). They found that among young social groups (formed for three years or less) both numbers of helpers and adult males were positively correlated with the number of surviving infants, but that in established groups (formed for over three years), only the number of helpers correlated with GRO (Bales et al. 2000).

Other than these correlational analyses, only one published study has attempted to model the predictors of GRO in callitrichids to date. Bales et al. (2001) modelled the effects of several maternal factors on female reproductive output in a population of *L. rosalia* that has two birthing seasons per year. Their analysis accounted for female identity, prior female reproductive success, repeat sampling of females across multiple years, and age. Female body mass predicted female reproductive output for litters in the first birth season, whereas that the number of helpers (as defined in Bales et al. 2000) explained offspring numbers in the second birth season. They concluded that mothers with increased helpers may carry infants less and thus be in better body condition for the subsequent birth season (Bales et al. 2001).

Prior analytical approaches have predominantly emphasized the role of adult males in driving GRO; yet there are substantial complications that should be addressed. Strict correlational analyses of mean GRO with numbers of individuals in different age-sex classes (Garber 1997; Koenig 1995) do not account for variable group sizes, which are often uneven across a species. For example, an assessment of data from a thirteen-year study on *Saguinus weddelli* (Goldizen et al. 1996)with groups containing 1-4 adult males revealed that 25% (12/47) of groups had only one male, 68 % (32/47) of groups had 2 adult males, while only 5 % (2/47) had 3 males, and 2 % (1/47) had four males – disparate sample sizes that preclude using means to test the effect of age-sex class on GRO as per Garber (1997). Koenig (1995) attempted to assess the impacts of group size on GRO across multiple studies, but these analyses did not consider uneven sampling or random variation between studies. Garber’s study (1997) observed a translocated population of wild-caught tamarins to Padre Isla that existed in isolation from many predators and faced reduced inter- and intraspecific competition, with the entire population consisting of only newly formed groups according to Bales et al’s (2000) criteria. Bales (2001) examined only the effects of female-factors on GRO in *L. rosalia*, excluding the potential influences of individuals from other age-sex classes (particularly adult males) from their model.

Most importantly, no prior analysis of callitrichid GRO has accounted for the variation in the reproductive capabilities, or developmental class, of individuals within the same age-sex class. Reproductive suppression of female callitrichids has been documented in both wild and captive settings (Barrett et al. 1990; Beehner and Lu 2013; Ziegler et al. 1987), and evidence of male dominance has been observed in moustached tamarins (*Saguinus mystax*), indicated by notable within-group differences (up to 53%) in testicular volumes (Garber et al. 1996). As Garber (1997) proposed, not all adults contribute equally to GRO in cooperatively breeding species. A distinction should be made between the set of individuals that copulate within a group (the mating system) and the smaller subset of individuals that contribute towards the gene pool of viable offspring (the breeding system) (Garber 1997). Kappeler and van Schaik (2002) refer to these as the social mating system and the genetic mating system respectively. Distinguishing between the two groups is dependent on identifying biological parents for all offspring, typically possible only via genetic analyses. Since most callitrichid groups contain a single male and female who make up the genetic mating system, with the rare exception of multiple males fathering offspring in the same litter (Díaz-Muñoz 2011; Huck, Löttker, Böhle, et al. 2005; Suárez 2007), we contend that an individual’s reproductive development, indicative of its potential to contribute to the gene pool, is most relevant to understanding the factors that drive reproductive output in primate groups with cooperatively breeding systems. Thus, our approach is concerned with the social, and not genetic, mating system.

Reproductive development has been assessed before through endocrine studies of derivatives of testosterone, estradiol, and prolactin among callitrichids in captivity (Ziegler et al. 1993) and in the wild (French et al. 2003; Löttker et al. 2004). However, wild studies are invariably challenged by the inability to collect blood for peptide hormones or adequate numbers of fecal steroid samples from known individuals across multiple ovarian cycles and breeding seasons (Löttker et al. 2004). One study that measured testosterone levels among wild moustached tamarins (*Saguinus mystax*) found that concentrations varied too widely during maturation to reliably determine an individual’s level of reproductive development, including between twin siblings (Huck, Löttker, Heymann, et al. 2005).

Another means to evaluate reproductive development is to assess dominance status in a group, the definitions for which differ by sex. Reproductive dominance has been ascribed to a single female in a group through endocrine monitoring, interactions with breeding males (marmosets: Sousa et al. 2005), and through age effects (i.e. the oldest female is the breeding female) (moustached tamarins: Garber 1997). Among males, this is not feasible in some cases (Huck, Löttker, Heymann, et al. 2005), but agonistic interactions have been used by some to identify a dominant male (Baker et al. 1993). Nevertheless, given observations of behavioral and physiological reproductive suppression among callitrichids, these metrics can fail to differentiate between individuals of varying reproductive potential, particularly in light of species and site-specific variation. Thus, since all individuals in an age-sex class cannot be assumed to possess similar reproductive potential, it is critical that developmental class, and not only age-classes, be assessed for possible impacts on GRO.

In this paper we compiled two datasets to test the impacts of various classes of individuals on GRO, while improving upon past analytical approaches. The first is a historical dataset of all published studies on wild callitrichids that provide data on the numbers of individuals in each age-sex class and the numbers of surviving offspring per year. We conducted a meta-analysis of this historical dataset that estimates the average magnitude of correlations between specific age-sex classes and GRO to test whether the effect is statistically significant from zero, while controlling for sample size differences across studies (Scheiner and Gurevitch 2001). Based on prior studies (Baker et al. 1993; Bales et al. 2000; 2001; Garber 1997; Koenig 1995) we predict that the number of adult males should be significantly correlated to GRO across studies.

The second dataset, from a 6-year study on saddleback (*Leontocebus weddelli*) and emperor tamarins (*Saguinus imperator*) at Los Amigos in southeastern Peru, consists of group compositions by sex-based developmental classes and the numbers of surviving offspring each year. We developed a novel method that uses multiple morphological variables collected via a mark-recapture program that reliably assigns individuals to one of three developmental classes – infantile, immature, and mature - that reflects their potential to participate in the social mating system of a group. We use these data to answer two questions on the factors that drive GRO. First, which developmental class has a significant effect on determining if a group has any offspring at all? To answer this question, we constructed a mixed effect logistic regression model with a binomial response variable (presence or absence of offspring), using males and females of different developmental classes as explanatory variables. Second, which developmental class has a significant effect on predicting the number of surviving offspring (0 to 3)? For this analysis, we constructed a generalized mixed effect model with number of offspring present as a discrete numerical response variable and sex-based developmental class compositions as explanatory variables. Finally, to control for the effects of group size on reproductive output, we used the proportions, and not absolute numbers, of individuals belonging to different developmental classes. For both questions, in line with prior studies, we predicted that mature males would have a significant effect on group reproductive output.

## Methods

### (a) Study Site and Subjects

We studied 21 groups of free-ranging saddleback tamarins (*Leontocebus weddelli*, formerly *Saguinus fuscicollis weddelli* (Buckner et al. 2015; Matauschek et al. 2010)) and emperor tamarins (*S. imperator*) at the Estación Biológica Río Los Amigos (EBLA) (12°34’S 70°05’W) in the Madre de Dios Department of southeast Perú annually across 6 seasons (2010-2015). We used a mark-recapture program (detailed protocol in (Watsa et al. 2015)) of 166 animals (106 *Leontocebus weddelli* and 60 *Saguinus imperator*) with an average of ~55 captures per year, for a total of 331 capture instances. At capture, infants were 4 to 7 months old, readily identifiable by facial pelage and dentition. The dominant breeding female in each group was fitted with a radio collar to facilitate tracking as a part of a larger behavioral study. Groups were followed for an average of 425 hours (range: 116 to 1135 hours) each season (May to August) and instances of mating and dispersal were recorded *ad libitum*. All groups were censused at least twice a month for group composition. In total, we monitored the study groups for a total of 2127 hours across the 6-year period. We recorded a total of 143 instances of mating across 33 males of both species.

### (b) Assigning developmental status

In this paper, we call those individuals that participated in mating, and who have the potential to contribute to the gene pool, as mature females or males. We were able to classify a female as mature if the female displayed a nipple length of > 3 mm for *Leontocebus weddelli* or > 4 mm for *Saguinus imperator* (Soini and de Soini 1990; Watsa 2013), indicating a prior birth record, regardless of whether multiple adult females or infants were present in the group. Mature males were considered to be any males that we observed copulating. We identified immature individuals in groups as those who were between 1 and 2 years of age i.e. were known to have been born in the prior census year. Thus, during a census, groups could consist of mature or immature members of both sexes, as well as any offspring born in that same year who would belong to the infantile class. Based on these criteria, we identified a subset of individuals of known reproductive developmental status that could be used to validate our models to predict reproductive developmental status in other animals.

During capture, we recorded length and width of genitalia and suprapubic glands to formulate indices of developmental status as follows: vulvar index (length + width), suprapubic gland area (length * width), average nipple length, and testicular volume (a semi-spherical estimate) (Garber et al. 1996; Soini and de Soini 1990). In 2.4 % (8/331) of captures, a measurement (not always the same one) was not recorded by accident. We included these 8 instances by replacing the missing values with the mean value for the measurement in that developmental class (if known, N=4), or in that age class instead (N=4). We mean-centred and scaled all measurements and indices by standard deviation for use in a principal components analysis by species-sex groups (Principal Components Analysis: FactoMiner package in R (Beehner and Lu 2013)). We used individual coordinate values from the first two principal components in a linear discriminant function analysis (LDA) to model assign individuals of unknown developmental status to three categories: mature, immature, and infantile. Resampling of individuals occurred 1 to 4 times per animal, with 51.8 % captured at least twice. To avoid pseudoreplication, we used mean individual component scores across years for animals with known developmental status to train the LDA functions. We checked each species-sex class for normality (q-q normal plots), linear relationships (linear regression), and homoscedasticity between developmental categories (Bartlett’s test of homogeneity of variance, *p* > 0.05). We omitted infant males of both species from the LDA due to limited variance causing heteroscedasticity; but since they were < 7 months old, this exclusion had no impact on adult and sub-adult male classifications. We calculated the percentage of known individuals that were correctly classified by this PCA-LDA model (Table 1), and used a MANOVA (manova: MASS package in R (Venables and Ripley 2002)) to test the null hypothesis that all predicted developmental status groups were indistinguishable based on individual component scores. All statistical analyses were performed in R v.3.2.2 (R Development Core Team 2015).

**Table 1:**
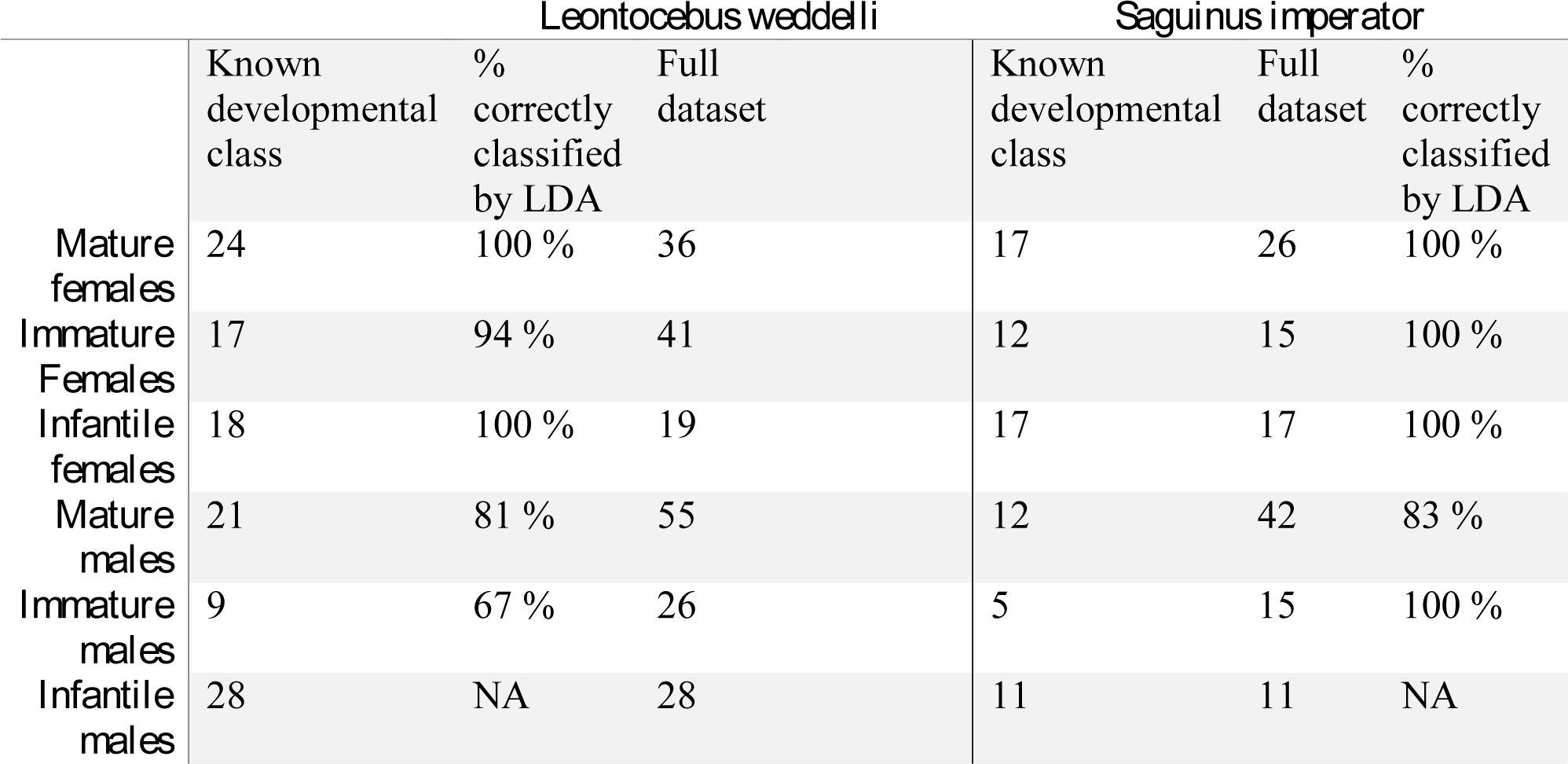
Sample sizes of developmental classes before and after the LDA model

### (c) Group reproductive output in the historical dataset

To assess the reliability of previous findings that numbers of adult males is strongly associated with GRO, we performed a meta-analysis of all available historical data. We utilized Google Scholar and Scopus to conduct a literature review for published information demographics and group reproductive output in wild callitrichid populations. We compiled a historical dataset from 15 studies published from 1976 to 2015 on wild populations of *Saguinus* spp. (including *S. geoffroyi, S. mystax, S. weddelli, S. tripartitus,* and *S. oedipus*)*, Leontopithecus caissara,* and *Callithrix jacchus*. Studies were included in our analysis only if they reported raw numbers of individuals per age-sex class and GRO across a minimum of 5 group-years (Appendix 2 in Electronic Supplementary Material). For the meta-analysis of numbers of adult males to GRO, we included an additional study (now N=16) on *L. rosalia* by Bales et al. (2000) by calculating the effect size from the sample size and Spearman’s rank correlations presented in the study. To combine data from multiple studies, we used a Spearman’s rank correlation weighted by the number of group-years in the study as a standardized effect size. In this dataset, social groups (within a study) and species (across multiple studies) were subject to repeated sampling over time, which could render certain data points non-independent. To control for interspecific differences, we added species as a moderator variable in a mixed effect meta-analysis of the historical dataset. Species did not have a significant effect and was subsequently removed; we proceeded with a random effects meta-analysis that does not assume equal effect sizes across studies. Regarding repeated sampling of a subset of groups in long-term studies, we feel that their inclusion does not bias our study more than their exclusion, which would drastically shrink the dataset. However, we use a more conservative significance level of p < 0.01 for the meta-analyses (see Gurevitch et al. 1992; Poulin 1994 for detailed explanation of this reasoning).

### (d) Group reproductive output in the Los Amigos dataset

Correlations are pair-wise, not predictive, and cannot control for group identity or species (Bolker et al. 2009). Further, group size can be controlled for by using proportions of individuals in each age-sex or developmental class. With this in mind, we first constructed a mixed-effect logistic regression model with a binomial error structure and a logit link function to predict a binary response variable - offspring presence or absence based on proportions of individuals of each developmental class as fixed factors. As per Bales et al. (2001) we also built generalised linear mixed models (GLMMs: lme4 in R (Bates et al. 2014)) with a Poisson error structure, response variable GRO (ranging from 0-3), and proportions of individuals per developmental class as fixed factors. We used saturated fixed-effect models to optimise random structures, and incorporated group identity, species, and year as needed to ensure independence of data points across all models. Correlation analyses were conducted on all pairwise combinations of explanatory variables and any fixed factor redundancies were removed. Each explanatory variable was plotted against the response variable to ensure that there were no nonlinear relationships. We established minimal models using Akaike Information Criterion (Akaike 1994) by backwards non-significant term deletion, retaining terms only if they reduced criteria by two units (Moreno et al. 2013). Minimal models were confirmed by performing a likelihood ratio test, which compares the difference in log-likelihoods of nested models with a Chi-square distribution. The residuals of best fit models were plotted to ensure that they were randomly distributed around zero.

### (d) Ethical Note

This study follows the Animal Behaviour Society Guidelines (Rollin and Kessel 2006) and American Society of Mammalogists’ Guidelines on wild mammals in research (Sikes and Gannon 2011). The study is part of an ongoing, long-term annual capture-and-release program that began at this site in 2009. In brief, we captured entire groups at baited compartment traps to which they are habituated, and processed and released them on the same day to minimize disruption and discomfort to the animals. Although based on previous capture protocols established for callitrichids (Savage et al. 1993), our study utilizes a novel two-step chemical restraint method that has improved recapture rates, virtually eliminated capture-related injuries, and has no visible effect on habituation for subsequent behavioral research (see Watsa et al. 2015 for protocol comparisons).

The Peruvian Ministry of the Environment (SERFOR) granted annual research and collection permits, and the Animal Studies Committees of Washington University in St. Louis and the University of Missouri St. Louis approved protocols.

## Results

### (a) Mean Group Reproductive Output per Age-Sex Class

As observed in one of the best longitudinal datasets on wild callitrichids (Cocha Cashu, *S. fuscicollis*) (Goldizen et al. 1996), the pattern of using average GRO that does not account for uneven sample sizes held true for the historical dataset. Disparate sample sizes per age-sex class resulted in overlapping 95% confidence intervals (eg. mean offspring =1.10 ± SD 0.87, CI: 0.94-1.27 in groups with two adult males while mean offspring = 0.93 ± SD 0.77, CI: 0.72-1.14 in groups with one adult male) (Figure 1). This precluded the use of mean GRO to evaluate the effect of age-sex class on GRO as per Garber (1997).

**Fig. 1.**
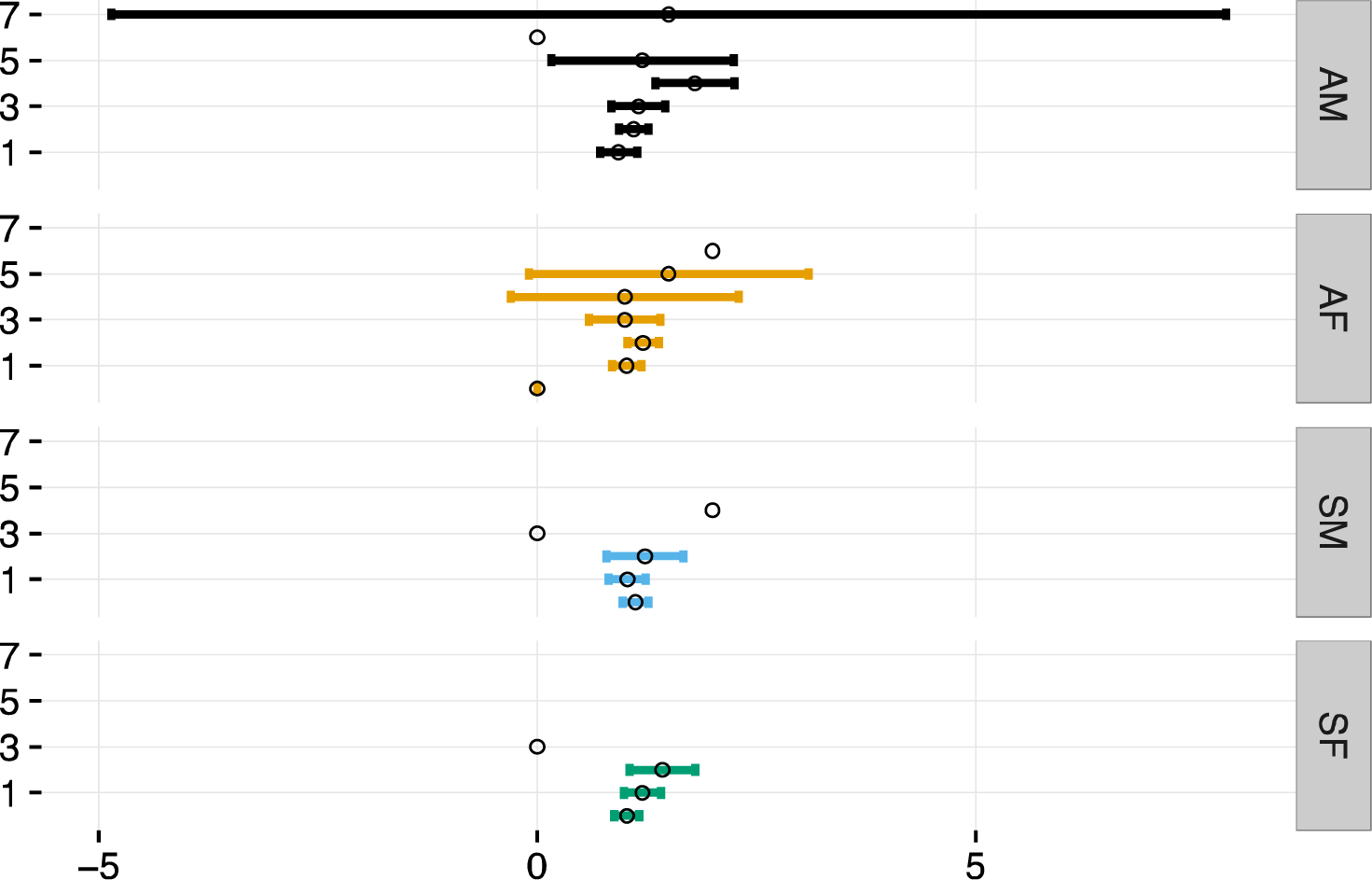
Average number of dependents (circles), with 95% C.I. (lines) depending on the number individuals from each age-sex class in the complete historical data set; adult males (AM), adult females (AF), sub-adult males (SM), sub-adult females (SF).

### (b) Meta-analyses of GRO in the Historical Dataset

A random-effects meta-analysis combining data from prior studies and the present study revealed significant Spearman’s rank correlations between adult females and GRO (weighted average rs (16) = 0.231, P < 0.001), as well as group size and GRO (weighted average rs (16) = 0.296, P < 0.001) (Figure 2). Adult males and subadults of either sex were not significantly correlated with GRO across studies (P > 0.05). These results remain unchanged when our study was excluded from the analysis.

**Fig. 2.**
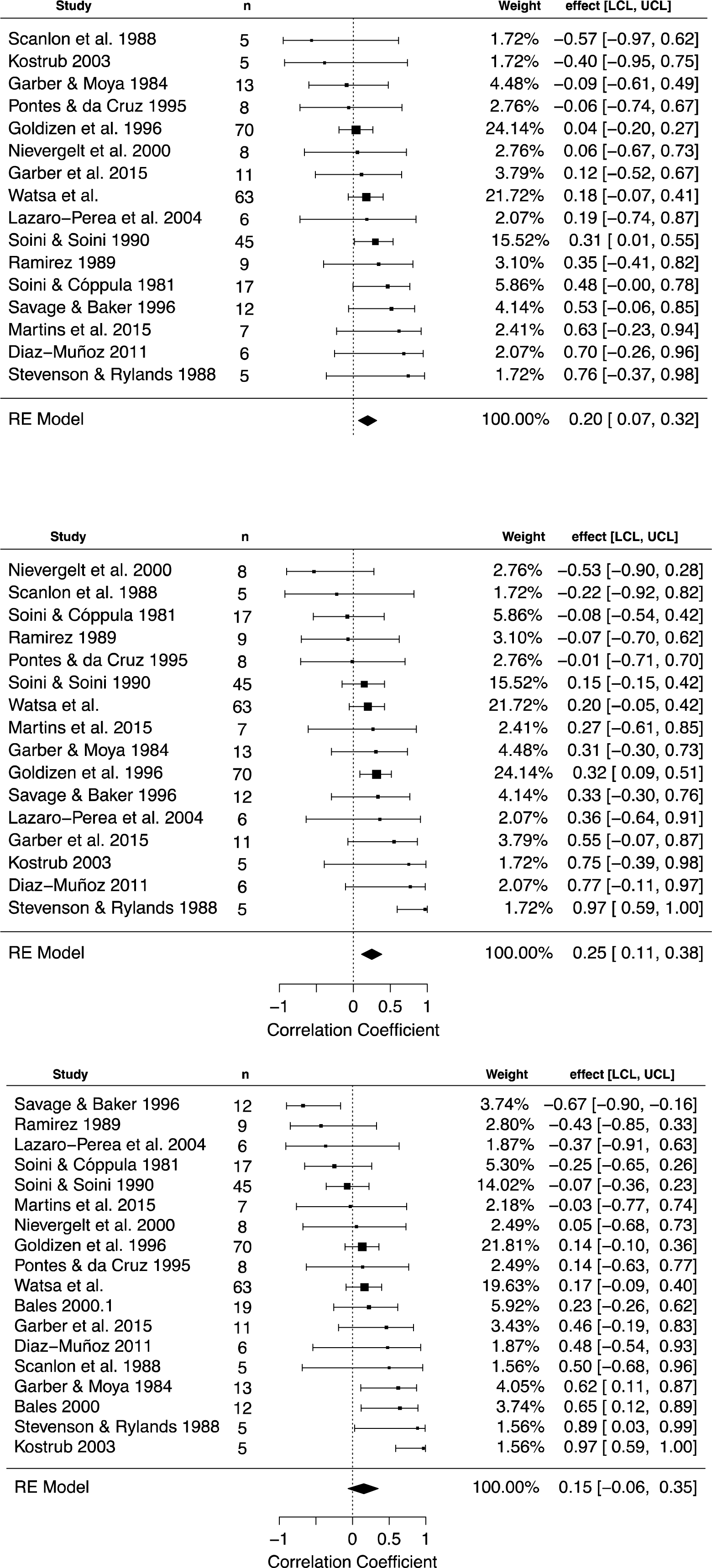
Forest plot showing the meta-analysis of a correlation of group reproductive output (GRO) with A) number of adult females (P < 0.001, N= 15); B) group size (P <0.001, N=15) and C) number of adult males (P > 0.05, N=16). Confidence intervals that do not overlap zero are generally not considered to be significant. Adult females (A) and group size (B) are significantly positively correlated with GRO across studies, while adult males (B) are not. These results are not altered when this study (Watsa et al.) is removed from the dataset.

### (c) Group Demographics from Our Study Population

Over 6 years we observed 21 groups across 63 group-years during which they could have reproduced, including 14 groups of *Leontocebus weddelli* sampled for a mean of 2.86 ± SD 1.35 years and 7 groups of *Saguinus imperator* sampled for a mean of 3.43 ± SD 1.27 years. Mean group sizes (Table 2), adult group sex ratios (males:females) (*L. weddelli*: 1.23 ± SD 0.63; *S. imperator*: 1.65 ± SD 1.34), and GROs (*L. weddelli*: 1.03 ± SD 0.87; *S. imperator*: 0.92 ± SD 0.88) were not significantly different between species (Welch’s Two Sample t-test, *p* >0.05). Across the study, 8.7% of all captured animals were infants, with 1-2 offspring per group and only one instance of three offspring. We also observed 7 instances of two mature females present in a single group – four cases in *L. weddelli* and three in *S. imperator*.

**Table 2:** Group compositions based on developmental class status. All figures are provided as mean number of individuals ± standard deviation (range).

| Species | N | Group Size | MF | ImF | MM | ImM | All Juvs | All Males | All Females |
| --- | --- | --- | --- | --- | --- | --- | --- | --- | --- |
| *Leontocebus weddelli* | 14 | 4.95 ± 1.63 (3-8) | 0.95 ± 0.50 (0-2) | 0.90 ± 0.78 (0-3) | 1.40 ± 0.98 (0-3) | 0.65 ± 0.74 (0-2) | 1.03 ± 0.86 (0-3) | 2.05 ± 0.90 (0-5) | 1.88 ± 0.69 (1-4) |
| *Saguinus imperator* | 7 | 5.21 ± 1.41 (3-8) | 1.08 ± 0.41 (0-2) | 0.67 ± 0.96 (0-3) | 1.71 ± 1.23 (0-4) | 0.63 ± 0.77 (0-2) | 0.92 ± 0.88 (0-2) | 2.33 ± 1.20 (0-6) | 1.96 ± 1.00 (1-4) |

Note: N = Number of unique groups.

### (d) The Developmental Class Model

In our model, the minimum requirements for factor analyses were satisfied, with an average of 19 and 23 samples per variable for the females and males, respectively. The first two dimensions represented an average of 86 % (range: 82 – 90 %) of total group variation. For all species-sex classes, Principal Components Analysis dimension 1 was determined by all morphological variables and Principal Components Analysis dimension 2 was determined primarily by nipple length in females and suprapubic area and body mass in males (Tables 1 and 2 in Appendix 1 of Electronic Supplementary Materials).

For animals with known developmental class (57.1 % of *Leontocebus weddelli* and 59.5 % of *Saguinus imperator*), the LDA correctly assigned 98.3 % of female *L. weddelli*, 100 % of female *S. imperator*, 76.7 % of male *L. weddelli*, and 88.2 % of male *S. imperator* (Figure 3, Table 1). The LDA classification mismatched one immature female to the infantile class (*L. weddelli*); four mature males were reclassified as immature males, three immature males as mature males (*L. weddelli*); and two mature males as immature males (*S. imperator*). The LDA successfully distinguished between developmental classes for females and males of both species (MANOVA, P <0.0001, Table 3), and we calculated mean values and ranges of morphological variables per species-sex group (Table 4). For both species, we observed variation in developmental classes in all age-classes except among infants (Figure 4).

**Fig. 3.**
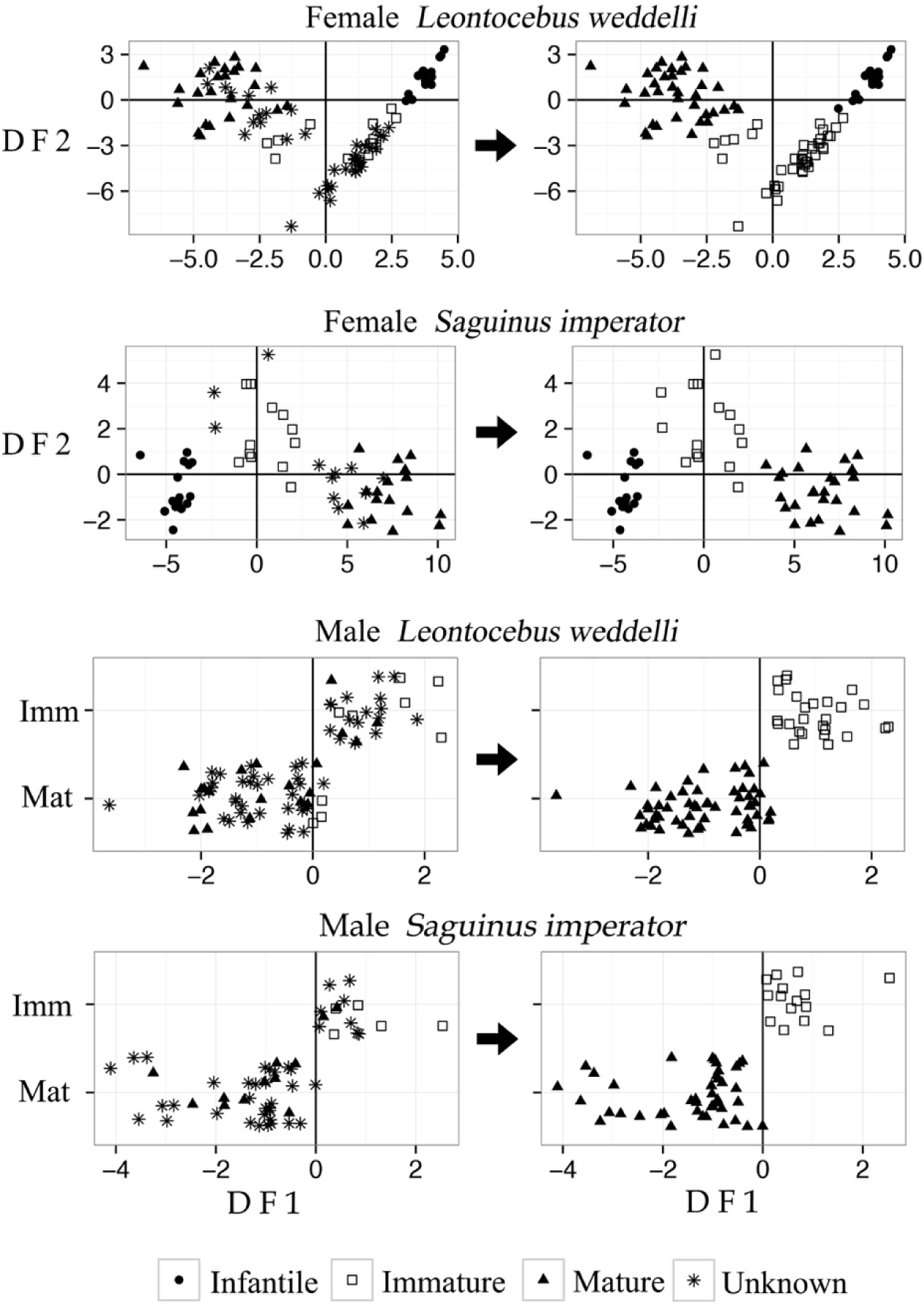
Developmental status by species and sex before (left) and after (right) implementing the PCA-DFA assignment model an classifying all individuals of uncertain status (star symbol) to a category based on reproductive morphology and weight. Female categories are differentiated by discriminant functions 1 and 2 (DF1 & DF2), while mature (Mat) and immature males (Imm) are differentiated by DF1 only; males in the infantile developmental class were removed from the DFA.

**Table 3:** MANOVA results distinguishing if developmental classes are significantly differentiated within all species-sex classes. Female assessment included three developmental classes (df=2), but males used only two (df=1).

| Species-Sex Class | Wilks' $\lambda$ | F | df | p-value |
| --- | --- | --- | --- | --- |
| Female *Leontocebus weddelli* | 0.0461 | 82.35 | 2 | <0.0001 |
| Female *Saguinus imperator* | 0.0348 | 56.67 | 2 | <0.0001 |
| Male *Leontocebus weddelli* | 0.3722 | 43.29 | 1 | <0.0001 |
| Male *Saguinus imperator* | 0.4756 | 19.48 | 1 | <0.0001 |

**Table 4:**
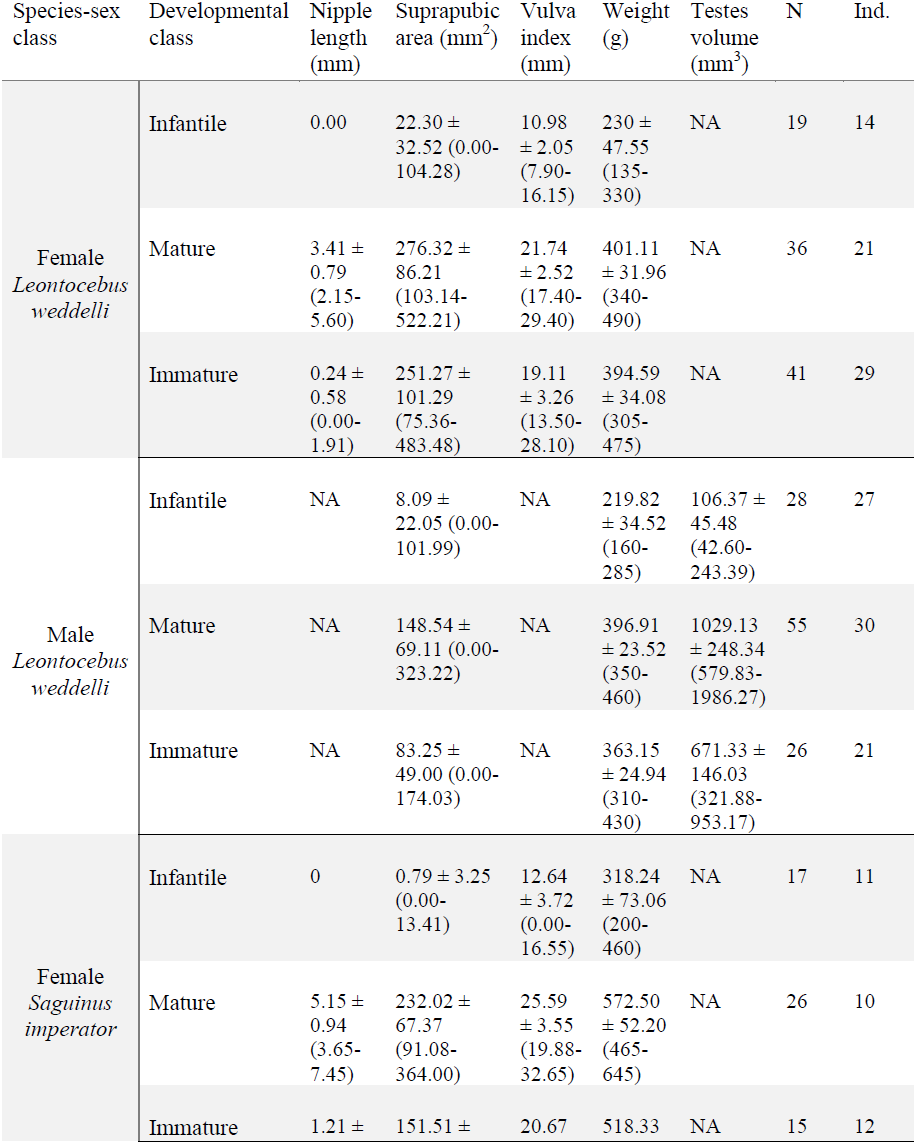
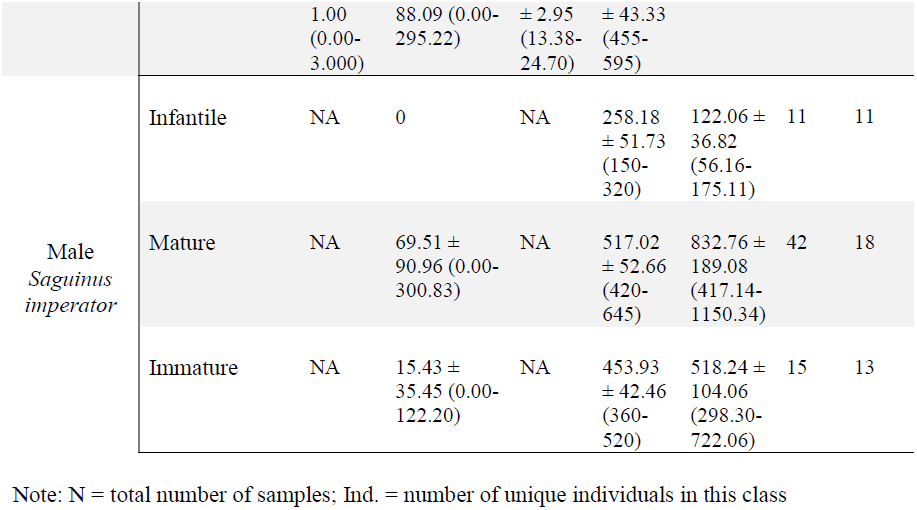
Morphological variables by developmental class. All values expressed as mean ± s.d. (range)

**Fig. 4.**
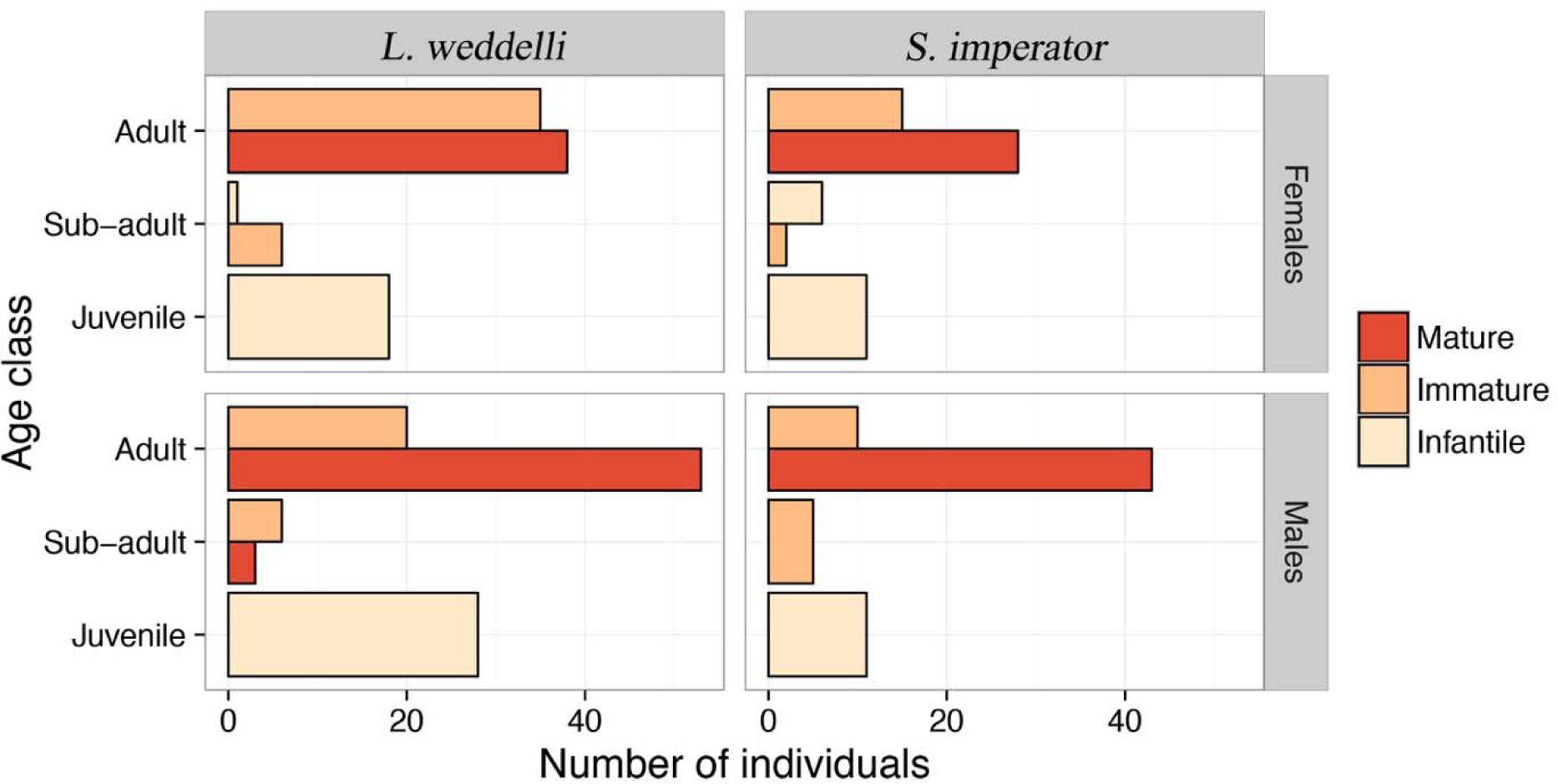
The distribution of developmental classes (mature, immature, and infantile) between age classes (adult, subadult and juvenile) for males and females of both callitrichine species at Los Amigos.

### (e) Group Reproductive Output in the Los Amigos Dataset

Our logistic regression model indicated that the proportion of mature males (B = 3.877, s.e. = 1.961, *χ*^2^ = 3.91, P < 0.05) was the sole significant factor in predicting the presence of offspring (Figure 5). The mean proportion of mature males in groups with no offspring (0.27 ± SD 0.23, N=22) was significantly lower than in groups with one or more offspring (0.41 ± SD 0.23, N=41; t=-2.32, df=43, P = 0.025). The proportions of mature females and immature males or females were not significant predictors of the presence of offspring in this analysis. However, a GLMM with offspring number as a discrete numerical response variable revealed that the proportion of mature females relative to group size (B = 3.559, s.e. = 0.962, *χ*^2^ = 13.69, P < 0.001) and group size (B = 0.343, s.e. = 0.128, *χ*^2^ = 7.15, P = 0.008) were the only two significant factors. Greater proportions of mature females and larger group sizes were significantly associated with GRO. The proportions of mature males and immature males or females were not significant predictors of GRO in the GLMM.

**Fig. 5.**
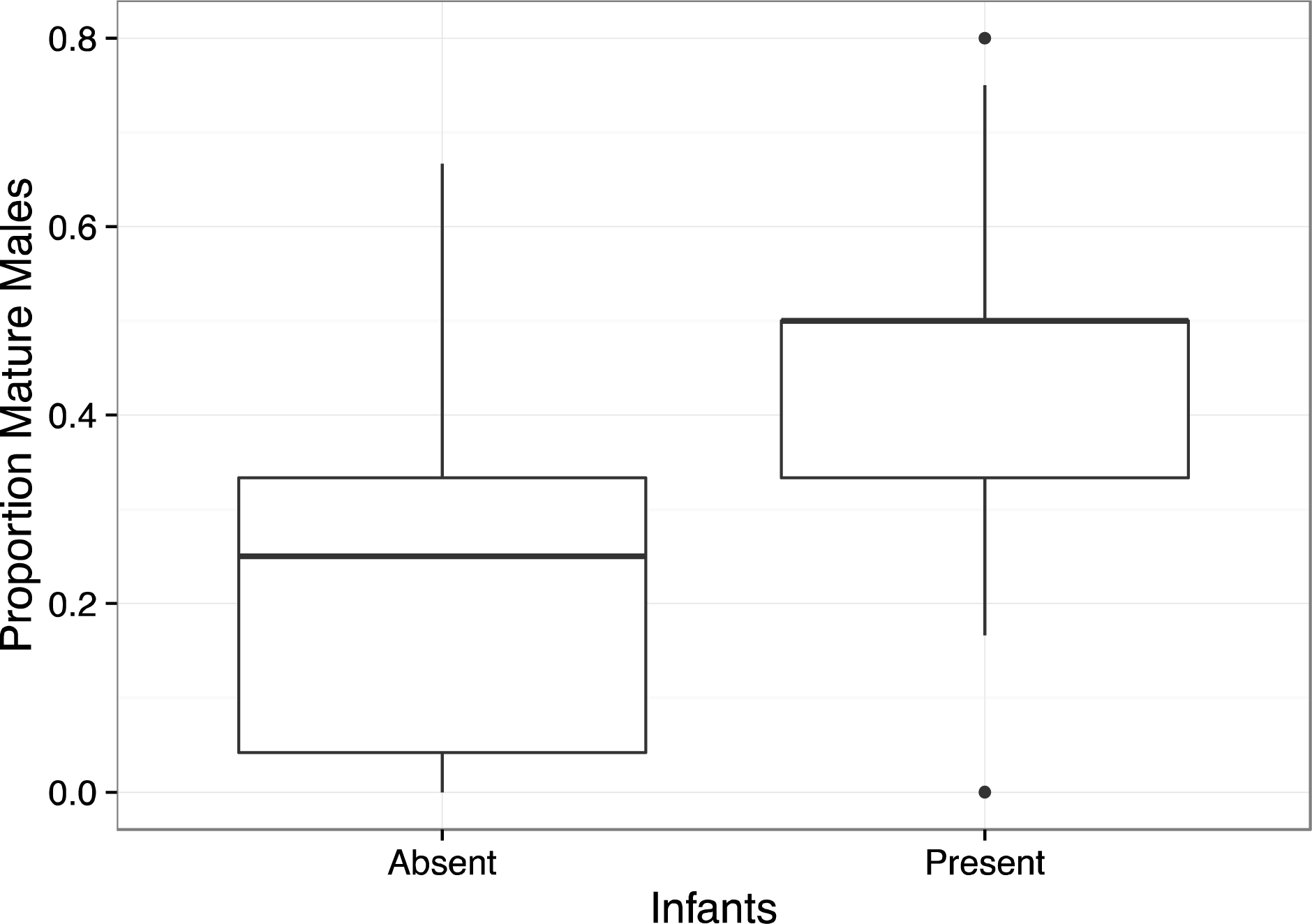
A box plot of the proportion of mature males in groups where infants are either present or absent. The two proportions are significantly different as revealed by a logistic regression model.

## Discussion

Like other callitrichids, both study species at Los Amigos twinned frequently and formed groups with multiple breeding females (Garber et al. 2015; Watsa 2013). Though these species diverged ~9.10-10.07 mya and are now placed in separate genera (Buckner et al. 2015; Matauschek et al. 2010), we noted no significant differences between them in mean group size, adult group sex ratios, or mean GRO.

### (a) Using Morphology to Determine Developmental Class

A method for reliably assigning developmental class is missing to date. Reproductive status has previously been evaluated in callitrichids through measurements of their genitalia (Soini and de Soini 1990). In addition, scent-gland morphology is known to signal oestrus, changes around parturition (*Callithrix jacchus* (Moreira et al. 2015)), and differs by sex (French and Cleveland 1984; Watsa 2013; Zeller et al. 1988); thus, it is likely related to developmental class (Watsa 2013). The proposed model utilised body weight and genitalia and scent gland morphology to assign animals into developmental classes. Females were more reliably assigned to the correct class than males, likely due to the availability of external validated measures in females, such as observed nursing and nipple lengths (Soini and de Soini 1990), which were missing for males. Higher resolution of male developmental class would require the inclusion of all or most copulation records, which was not feasible as copulation is cryptic among arboreal primates (Campbell 2006) and of short duration (1-12 s) in tamarins (Watsa 2013). Nevertheless, our model successfully discriminated between developmental categories for all species-sex classes, confirming that all animals of a particular age-sex class did not have equal reproductive capabilities.

### (b) Drivers of Group Reproductive Output

The potential causes for variability in the number of males in a primate group have been long debated (Carnes et al. 2011; Heymann 2000; Kappeler 2000; Ridley 1986). The number of adult males in a group have been proposed to increase with shorter breeding seasons (Ridley 1986), since a single male probably cannot successfully monopolize multiple reproductively synchronised females (Dunbar 2000). While some research supports this premise (Carnes et al. 2011), it has also been suggested that primate males simply “go where females are” (Altmann 1990). Cross-species analyses that control for phylogeny show that these theories are not necessarily exclusive - the number of males is tightly positively correlated with the number of females in primate groups across species (Mitani et al. 1996), but female breeding synchrony or overlap can predict adult male numbers after female numbers are controlled for (Nunn 1999). Other theories for larger numbers of males in groups include heightened predation risk (Savage et al. 1996; van Schaik and Hörstermann 1994) or, as with callitrichids, the necessity for alloparents due to the high costs of caring for twin infants (Heymann 2000; Tardif 1994; 1997). But among callitrichids, with the exception of *Callimico*, Heymann (2000) found the number of adult males to be positively correlated with litter mass gain and daily path length, implying that adult males are necessary to counter increased costs of infant care across several callitrichine genera. This conclusion was further supported in an extensive cross-genera analysis of the effect of infant care costs on variation in reproductive behaviors (Díaz-Muñoz 2015). For example, *Saguinus* was identified to exhibit the highest infant care costs of all callitrichine genera, which was correlated with high subordinate female reproductive suppression, natal individuals dispersing early, and increased offspring produced by the dominant female. Most importantly, this conclusion was partially supported among species with high (*Saguinus* spp.) and intermediate (*Callimico goeldii* and *Leontopithecus rosalia*) infant care costs due to the assumption that the number of adult males is indeed correlated with infant survival (Bales et al. 2000; Koenig 1995). However, our data challenge the validity of this assumption for all levels of GRO.

Meta-analyses form a robust methodology to summarize the outcomes of a range of research studies and have been used for some time in the field of medicine as a powerful tool for health-related decision-making (Gurevitch et al. 1992; Scheiner and Gurevitch 2001; Vetter et al. 2013). The meta-analysis presented here coalesced Spearman’s rank correlations of GRO with numbers of adult males and females and group size across 16 studies on three callitrichid genera. By using a random effects model, our analysis did not assume that different studies shared a common effect size, which is an unlikely assumption across ecological studies. Significantly, our findings did not support the expected correlation of numbers of adult males with GRO as we had predicted; instead, group size and the number of adult females were discovered to be significant drivers of GRO across studies. These results provide the first inkling that identifying drivers of GRO is more complex, and that the role that adult males play could be affected by other factors, such as sexual maturity and finer distinctions in definitions of GRO.

Focusing on the data from Los Amigos, we were able to show that mature males, of all males, had a significant effect not on the full range of outcomes of GRO, but on whether a group had any infants at all. Males did not explain the increase in offspring numbers from zero to two or more infants. Although prior studies have, for the most part, not identified females as playing a significant role in this aspect, the most comprehensive dataset on wild callitrichids (*L. rosalia* from Poço das Antas Reserve in Brazil) did indeed show that female factors are important to GRO (Bales et al. 2001). They found that female body mass was the most significant factor across birth peaks. We utilize body mass and other morphological characteristics in ascribing developmental class, and thus, our data supports this finding on *L. rosalia*.

Higher proportions of mature males in a group appear to be necessary across *Leontocebus weddelli* and *Saguinus imperator* in ensuring that groups have offspring at all. Beyond that, the sexual maturity and proportion of females in a group dictate the probability of having a higher number of offspring. Increased GRO as a onseqeunce of higher proportions of mature females could have been facilitated in a variety of ways in our dataset from Los Amigos. First, we observed a case of allonursing of infants by a mature female *L. weddelli* who most likely lost her own infants at birth, which permitted the twin infants to nurse for close to six months of age instead of being weaned at age three months, as is typical (full account in Watsa 2013). Second, we report a case of multiple breeding female *L. weddelli* in which we recorded a pair of infants that differed in age by approximately two months, based on their tooth eruption schedules. Finally, we also report an instance of observing three offspring in a single group of *L. weddelli*, of approximately the same age, implying that multiple breeding females had produced offspring that were raised successfully to weaning. Further, until now, there has been no way to assess disparities in reproductive output between equally sized groups. Using our analyses, we find that additional variation in GRO is explained by group size.

Diaz-Muñoz (2015) identified male-biased group composition as a trait shared by all callitrichine groups, while the number of males that mate and sire young varied across genera. This variability was tied closely to individual reproductive strategies that remain flexible in the context of differing infant care cost scenarios. Among callitrichids and some other primates, there is often a marked disconnect between age and reproductive capability (Barrett et al. 1990; Beehner and Lu 2013; Ziegler et al. 1987). These insights both reinforce and challenge certain perspectives on the proximate causes for cooperative breeding behaviour in callitrichids. Mature females, who habitually produce twins, benefit from the presence of alloparents chiefly in that they are able to reduce the high energy costs associated with carrying and provisioning infants, while reducing competition for resources for their infants with others (Fite et al. 2005; Tardif 1994). Mature males may be motivated by having access to mating opportunities with a dominant female, rather than seeking pair-bonds with subdominant, less experienced females. Males also ensure the success of offspring they may have fathered and lower overall energetic investment per infant (Achenbach and Snowdon 2002; Beehner and Lu 2013; Burkart 2015; Santos and Martins 2000; Washabaugh et al. 2002). Moreover, all mature individuals can gain improved territory defence in larger groups (Bales et al. 2000; Lazaro-Perea 2001).

However, the motivation for immature alloparents is less straightforward. The group augmentation hypothesis states that offspring survival will benefit alloparents by increasing the number of future helpers, thus reducing energetic demands on them in the future (Kingma et al. 2014). This explanation is not supported by our data that indicated that immature individuals of either sex did not influence GRO. Rather, our findings support the idea that maximum GRO could actually be hindered by the reproductive suppression of subordinate females (Bales et al. 2000), since groups with multiple mature females had greater GRO. It would be worthwhile to collect data on GRO from other callitrichid populations that have reported multiple breeding mature females in a single group (see reviews: Digby and Ferrari 1994; Hilário and Ferrari 2010; Smith et al. 2001), a variable we were unable to extract from the historical dataset compiled in this study. While our findings regarding immature helpers conflict with previous studies of callitrichids that found them to be integral to GRO (Bales et al. 2000; Clutton-Brock et al. 2001), they conform with what we know of other cooperative breeding species. In meerkats (*Suricata suricatta*), similar modelling approaches revealed that helpers do not have a direct effect on litter sizes at birth or pup weights at weaning, which were influenced by maternal weight instead (Russell et al. 2003). Among European badgers (*Meles meles*), the impact of helper numbers on GRO was actually mediated by territory quality (Woodroffe and Macdonald 2000), in line with our finding that immature individuals benefit GRO via increased group size. The next steps in understanding the roles of both biological and non-biological parents in influencing GRO should take into consideration within-group genetic relatedness to address the role of the genetic mating system, and detailed comparisons of alloparenting behaviors of immature individuals with that of mature individuals in the population at Los Amigos.

## Electronic Supplementary Material

Supporting Information on Principal Components Analysis (Appendix 1) and the historical dataset (Appendix 2) are available online.

## Conflict of Interest

The authors declare that they have no conflict of interest.

## Authors’ Contributions

GE and MW conceived of and designed the study. MW determined the initial breeding status models, collected field data and drafted the manuscript; GE collected field data, carried out the statistical analyses, and drafted the manuscript; ER collected field data and helped draft the manuscript. All authors gave final approval for publication.

## Funding Statement

This study was funded by Field Projects International, the American Society of Mammalogists, Idea Wild, the Animal Behavior Society, Lambda Alpha, the International Primatological Society, the American Society of Primatologists, the Society for Integrative and Comparative Biology, Sigma Xi, and Trans World Airlines.

## Acknowledgements

This study was funded by Field Projects International, the American Society of Mammalogists, Idea Wild, the Animal Behavior Society, Lambda Alpha, the International Primatological Society, the American Society of Primatologists, the Society for Integrative and Comparative Biology, Sigma Xi, and Trans World Airlines. We would like to acknowledge the logistical support of the Amazon Conservation Association, staff at EBLA, and the Ministry of Agriculture in Peru. In addition, we want to thank all wildlife handling research assistants who assisted in putting together this dataset with FPI and PrimatesPeru over the years. We would also like to acknowledge the comments by two anonymous reviewers and the editor of the International Journal of Primatology.

